# Heliox effect on expiratory flow in master athletes: a comprehensive analysis coupling experimental measures and computational modeling

**DOI:** 10.64898/2026.11.24.769050

**Authors:** Valentin Mons, Jonathan Stephano, Toni Haddad, Colin Lavigne, Olivier Meste, Grégory M. Blain, Benjamin Mauroy

**Author notes:** **Corresponding author** Benjamin Mauroy, Laboratoire J.A. Dieudonné, Université Côte d’Azur CNRS UMR 7351., Parc Valrose 06108, Nice, France.

## Abstract

**Introduction:** Expiratory flow is determined by airway geometry and inhaled gases’ properties. Maximal expiratory flow-volume (MEFV) curves decrease with aging that reduces airway caliber and increase resistance. However, the independent contributions of airway geometry and gas properties to expiratory flow remain difficult to quantify experimentally. We combined experimental measurements and mathematical modeling to evaluate these contributions in master athletes, a model of successful aging.

**Methods:** Spirometry was assessed in 14 master athletes (69.3±5.7yr) breathing air or heliox (HeO_2_). A lung model comprising 17 generations of compliant airways with symmetrical bifurcations was calibrated using measured MEFV curves obtained with air. The model simulated MEFV curves with HeO_2_, isolating the effects of gas properties from airway geometry. To mimic the effect of aging in our lung model, MEFV curves were simulated with 15% and 30% reductions in airway size. MEFV curve shape was characterized using the slope ratio method.

**Results:** Heliox increased expiratory flow at higher lung volumes, and slope ratios were similar between air and HeO_2_ in both experimental and simulated MEFV curves. Higher tracheal flow increased the generations at which HeO_2_ reduces expiratory resistance. Simulated reductions in airway caliber amplified the effect of heliox on expiratory flow.

**Conclusion:** Heliox-induced reductions in airway resistance did not alter MEFV curve shape, and our model elucidates how reduced airway caliber amplifies the flow response to heliox through changes in resistance distribution across airway generations. This provides a framework for investigating the contributions of airway geometry and gas properties to expiratory flow with aging.

## INTRODUCTION

Flow dynamics within the lungs are governed by airway geometry and the physical properties of inhaled gases. These dynamics shape the movement of air during everyday breathing and determine the volumes and flow rates measured to evaluate lung health. Spirometry, typically performed while breathing air, requires maximal inspiratory and expiratory efforts from which the maximal expiratory flow-volume (MEFV) curve is obtained (1, 2). Analysis of the MEFV curve is clinically important as it can identify, for example, pathological reductions in expiratory flow, such as those caused by airway obstruction (3).

The MEFV curve is altered by the age-related decline in lung elastic recoil, which leads to airway narrowing and increases airway resistance and turbulence (4, 5). These age-related changes make older adults more likely to reach the mechanical ceiling of expiratory flow, particularly during exercise, a phenomenon known as expiratory flow limitation. Reducing the density of the inhaled gas can help quantify the contribution of inertial and turbulent forces to resistance and expiratory flow limitation. This can be achieved by replacing nitrogen with helium using heliox (HeO_2_), a mixture of 79% helium and 21% oxygen, which reduces gas density and thereby modulates gas convection to the alveoli. The properties of heliox promote more homogeneous ventilation, particularly in smaller airways where increased gas velocity amplifies their resistance to flow (6). Heliox is commonly used clinically to alleviate the work of breathing in individuals with airway obstruction (7) and experimentally to facilitate expiratory flow during exercise (8, 9).

To isolate the effects of aging from those of physical inactivity, disease, or deconditioning, we studied master athletes as a model of successful aging (10–12). In this population, by reducing gas density and lowering air inertia, heliox can reveal to what extent expiratory flow depends on airway resistance. However, pulmonary function measurements alone cannot determine where along the bronchial tree these resistance-dependent effects arise or how they relate to age-related changes in airway structure. Advanced imaging could provide such structural information, but its use may be technically challenging or limited by patient safety concerns, including radiation exposure. Coupling experimental measurements with numerical modeling of the lungs may therefore provide a more comprehensive understanding of the physical mechanisms governing expiratory flow in the aging lung. Specifically, combining heliox testing with lung modeling in master athletes breathing heliox may help determine how aging-related airway narrowing and density-dependent effects interact to shape expiratory flow.

Previous studies have used numerical modeling to compare airflow distribution during breathing with air and heliox, and to investigate mechanisms underlying reductions in the work of breathing (13–16). These models typically emphasize the role of airway geometry in determining airflow distribution, for example, by illustrating how disease-related reductions in airway caliber can increase airway resistance and impair expiratory flow. Among the different approaches used to represent airway geometry, the pulmonary tree model developed by Lambert (17) is based on the idealized lung geometry proposed by Weibel (18). This model represents the conductive airway tree using a cascade of 16 successive symmetrical bifurcations of cylindrical airways originating from the trachea. In this model, all airways within a single generation are identical, which allows to reduce the system to a series of 17 cylindrical branches. In this framework, flow distribution depends mainly on tracheal flow, surrounding tissue pressures, and hydrodynamic resistances. By adjusting gas properties such as density and viscosity within this model, we can predict flow dynamics and resistance distribution throughout the airways, thereby identifying regions where flow limitation or airway constriction may occur. Investigating the response to heliox in master athletes may therefore provide mechanistic insight into how flow inertia influences expiratory flow and airway resistance in the aging lung.

Accordingly, the present study evaluated whether reducing density-dependent airway resistance with heliox improves expiratory flow in master athletes. By combining experimental measurements with a compliant airway tree model, we aimed to determine how gas properties and airway geometry interact to shape flow patterns, resistance distribution, and potential sites of expiratory flow limitation in the aging lung.

## MODEL AND METHODS

### Subjects’ characteristics

Fourteen master athletes (2 females, 12 males) participated in this study. Participants were eligible if they were older than 60 years and had a 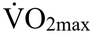 exceeding 130% of age-matched predicted values (19). All participants were healthy, non-smokers, were not under any medication regimen, had no symptoms indicative of cardiovascular respiratory or neuromuscular disorders, did not have respiratory infection during the 6 weeks prior to participation and had normal pulmonary function (**Table 1**). Prior to data collection, written informed consent was obtained from the participants. All study procedures were approved by the local Human Research Ethics Committee and adhered to the Declaration of Helsinki.

**Table 1.** Participants’ characteristics, pulmonary function and peak exercise data.

|  | **Measure**<br>**(n = 14)** |
|---|---|
| **Demographics** |  |
| Age (y) | 69 ± 6 |
| Height (cm) | 172 ± 8 |
| Weight (kg) | 67 ± 9 |
| BMI (kg/m<sup>2</sup>) | 22.5 ± 1.3 |
| **Pulmonary function** |  |
| FVC (%predicted) | 114 ± 14 |
| FEV<sub>1</sub> (%predicted) | 109 ± 14 |
| FEV<sub>1</sub>/FVC (%predicted) | 95 ± 10 |
| FEF<sub>75</sub> (%predicted) | 150 ± 46 |
| FEF<sub>25-75</sub> (%predicted) | 121 ± 39 |
| **Peak exercise** |  |
| Peak work rate (W) | 232 ± 47 |
| Relative peak work rate (W/kg) | 3.5 ± 0.5 |
| V̇O<sub>2max</sub> (L/min) | 3.32 ± 0.61 |
| V̇O<sub>2max</sub> (mL/min/kg) | 49 ± 7 |
| V̇O<sub>2max</sub> (%predicted) | 147 ± 14 |

All values are expressed as means ± SD. BMI, body mass index; FVC, forced vital capacity; FEV<sub>1</sub>: forced expiratory volume in one second; FEF<sub>75</sub>, forced expiratory flow at 75% of FVC; FEF<sub>25-75</sub>, forced expiratory flow between 25 and 75% of FVC; V̇O<sub>2</sub>, oxygen uptake.

### Experimental overview

Participants visited the laboratory on 3 different occasions. During the first session, participants were familiarized with experimental equipment and pulmonary function testing and performed a maximal and graded cycling test to exhaustion on a cycle ergometer to verify their physical fitness (Velotron, RacerMate, Seattle, WA, USA) as previously described (9). Participants underwent pulmonary function assessment at rest (see below), both before and after the cycling test, following the recommendations of the American Thoracic Society and European Respiratory Society (1). During the two other sessions, pulmonary function assessments were performed breathing either air (AIR, 79% nitrogen - 21% O_2_) or heliox (HeO_2_, 79% helium – 21% O_2_) following a randomized, single blind cross-over design.

### Pulmonary function testing

After receiving a thorough explanation of the procedures for pulmonary function testing, the participants were asked to perform 6-8 spirometry maneuvers. Participants were asked to perform a forceful and rapid blast out as hard and fast as possible from total lung capacity (TLC) to residual volume (RV), followed by a maximal and full inspiration from RV to TLC. Encouragement was provided to ensure that participants were making maximum efforts and that expiration lasted for at least 6 seconds (1, 2). These maneuvers aimed at measuring forced vital capacity (FVC), forced expiratory volume in 1 second (FEV_1_), FEV_1_/FVC ratio, peak expiratory flow (PEF), the volume of air breathed out at PEF (PEF_volume_), slope ratio (SR), and forced expiratory flow from 25 to 75% of vital capacity (i.e., FEF_25_, FEF_75_, FEF_25-75_). Throughout each attempt, participants received visual feedback of the flow-volume loops on a monitor.

### Data collection and processing

Expiratory flows were measured with one calibrated heated pneumotachograph connected to a two-way nonrebreathing valve (model 2730, Hans Rudolph, Kansas, USA). The pneumotachograph was calibrated with air or heliox depending on the experimental condition. Calibration was performed with a 3-L syringe (model 5530, Hans Rudolph, Kansas, USA), using three different flow rates (<70 L/min, 100-120 L/min and >150 L/min) and six different volumes (i.e., from 0.5 to 3 L) with the volume being calculated by numerical integration of the flow signal. During the whole session, raw flow and volume data were continuously recorded at a sample rate of 2000 Hz using a 16-channel analog-to-digital data acquisition system (Powerlab 16/35, AD Instrument, Australia). Data analysis was performed using MATLAB (MathWorks Inc., USA).

### MEFV curvilinearity analysis

Slope ratio was calculated to quantify the shape of the MEFV curves. A tangency and a chord line were drawn at points of interest calculated in 2% increments between 20% and 80% of maximal volume following previously detailed methods (20, 21). This portion of the MEFV curve was chosen as it corresponds to the effort-independent portion of the MEFV curve and expiratory flow cannot be enhanced by an increase in alveolar pressure. Values close to peak expiratory flow and residual volume were not selected to avoid effort dependent values (22). The tangent line was drawn using a volume difference of 200 mL above and below the point of interest and the chord line corresponded to the line intersecting the point of interest and the end of the FVC. The average SR value of each point of interest over the analyzed range of FVC were reported. A SR value of 1 corresponds to a linear MEFV curve, a value <1 indicates smaller change in flow per volume unit corresponding to a convex shape and a value >1 indicates greater change in flow per volume unit corresponding to a concave (20, 21).

### Model

The mathematical model was used to determine whether the expiratory flow response to heliox could be explained by changes in gas properties alone or whether age-related airway narrowing was also needed to reproduce the observed behavior. As the complexity of the human lung cannot be fully captured, the purpose of computational modeling is not to reproduce the experimentally measured MEFV curves exactly, but rather to provide a mechanistic framework for understanding how specific physical and structural determinants influence expiratory flow by studying mechanisms that cannot be isolated experimentally in vivo. Therefore, the model is intended to provide mechanistic insight and allows airway caliber to be modified independently of other pulmonary mechanisms, revealing how airway narrowing amplifies the expiratory flow response to reduced gas density. This approach helps explain the interaction between airway geometry and gas properties beyond what can be inferred from spirometry alone. Experimentally measured MEFV curves obtained with air were used as model inputs to simulate corresponding MEFV curves under HeO_2_ conditions to determine where along the bronchial tree the resistance-dependent effects arise (**Figure 1**). Expiratory volume was calculated by numerical integration of the flow signal. We then applied the model proposed by Stéphano and Mauroy (23) to estimate flow hydrodynamics and bronchial mechanics throughout the airway tree. By modifying gas density and viscosity within the model, we estimated how heliox influences flow distribution and resistance across airway generations. To isolate the contribution of age-related airway narrowing, heliox simulations were performed under three airway geometry conditions: unchanged lumen area (SIMU, control/younger adult condition), 15% lumen area reduction (SIMU15) and 30% lumen area reduction (SIMU30) (4, 24).

**Figure 1.**
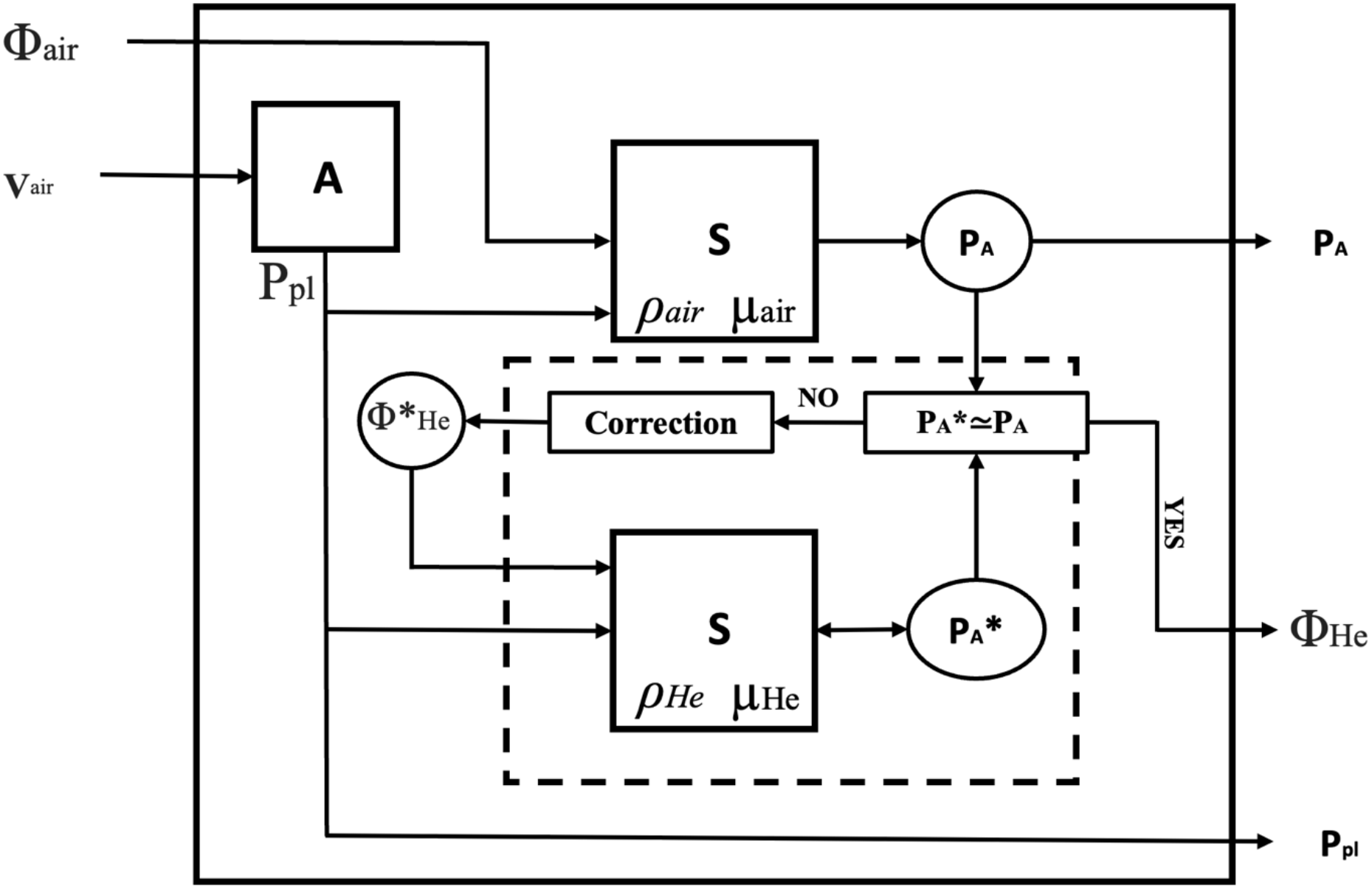
Mathematical model used to simulate maximal expiratory flow-volume curves curve with heliox. Φ_air_, expiratory flow with air; Φ_He_, expiratory flow with heliox;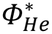, potential expiratory flow with heliox; V_air_, volume with air; ρ_air_, air density; ρ_He_, heliox density; µ_air_, air viscosity; µ_He_, heliox viscosity; P_pl_, pleural pressure; P_A_, alveolar pressure with air; 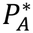, alveolar pressure with heliox; **A**, Agostoni and Rahn model; **S**, Stephano model (23); circled elements, numerical variables, squared elements, numerical procedure.

#### Lung airflow model

This model predicts the pressure distribution from the total flow rate *Φ*_*0*_ and the pleural pressure *P*_*pl*_ in an airway network which mimics the human pulmonary tree. The conductive bronchial tree is modeled as an airway tree with symmetrical bifurcations and 17 generations of airways. Each branch of this tree is assumed cylindrical and fully defined by a length and a cross-sectional area, as proposed by Lambert et al. (17). The Lambert et al. model characterizes the airway mechanics by a relationship between airway cross-sectional area and the pressure difference between the inside and the outside of the airways. In our context, the outside pressure is considered equal to the pleural pressure *P*_*pl*_.

Within each branch of a generation *n*, considering Hagen-Poiseuille law and the assumption that there is no fluid acceleration in the airways, a pressure drop *q*_*n*_ occurs, depending on both flow rate *Φ*_*n*_ and hydrodynamic resistance *R*_*n*_ of the branch:

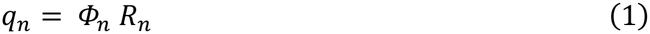

For steady laminar flows, mainly in distal airways, hydrodynamic resistance is determined using Poiseuille equation and depends on airway length *L*_*n*_, airway surface area *A*_*n*_ and the viscosity of the fluid *µ*.

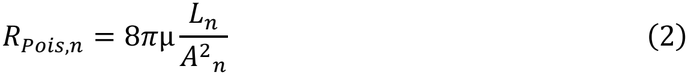

However, when higher flow rates are involved, mainly in proximal airways, the flow profile is deformed by the inertial effects and geometry. Thus, the hydrodynamic resistance of the fluid increases and becomes greater than Poiseuille resistance. In this context, we used the resistance factor *Z*_*n*_ proposed previously (25) to mimic the increase of hydrodynamic resistance with inertia effects:

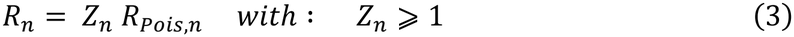

The resistance factor *Z*_*n*_ is superior to 1 for non-Poiseuille flow and depends on Reynolds number, 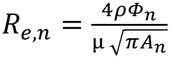, with ρ corresponding to gas density.

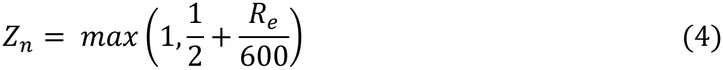

Here, we see that if the Reynolds number is inferior to 300, the flow is considered as a Poiseuille flow, and the factor *Z*_*n*_ is equal to 1. Additionally, in our model, flow is always considered as laminar and turbulence effects (*R*_*e,n*_> 2000) are neglected (15).

As illustrated by Reynolds number, the viscosity of the gas and especially its density play a major role in modulating resistance factor when inertia effects are involved. In Stephano and Mauroy’s study, the hydrodynamic resistance *R*_*n*_ is expressed in a sum of one term related to the gas viscosity *m* and a second term related to the gas density ρ (23), that includes the contribution of axial inertial effects to the hydrodynamic resistance.

Helium is an inert gas with a lower density and a slightly higher viscosity than nitrogen (ρ_He_ = 0.17 vs. ρ_N2_ = 1.17 g.m^-3^ and µ_He_ = 19.4×10^-6^ vs. µ_N2_ = 17.9×10^-6^ n.sec.m^-2^ at 37°C and 760 mmHg for helium and nitrogen, respectively) (26). Heliox and air therefore have different densities and viscosities (ρ_HeO2_ = 0.434 vs. ρ_air_ = 1.125 g.m^-3^ and µ_HeO2_ = 22.56×10^-6^ vs. µ_air_ = 18.95×10^-6^ n.sec.m^-2^ at 37°C and 760 mmHg for heliox and air, respectively) (16).

Breathing a low-density gas like heliox markedly reduces inertia effects and hydrodynamic resistance when expiratory flow rates are high. Comparing flow dynamics with heliox versus air enables to clarify the contribution of inertia on expiratory flow and reveals how age-related airway narrowing amplifies the effect of these resistances.

These relationships are applicable to a single airway and can be expanded to the lungs by including flow variations in each subsequent generation. Considering that each branch from a given generation has similar physical properties and that air and heliox are incompressible, the total flow rate in a generation is the product of the number of airways in that generation, i.e. 2^n^ with the air flow in an airway, *Φ*_*n*_. Then, using air flow rate in the trachea *Φ*_*0*_, we obtain *Φ*_*n*_ *= 2*^*-n*^ *Φ*_*0*_. Hence, only *Φ*_*0*_ is needed to determine all the flow distribution in the tree model. To summarize, the model predicts alveolar pressure *P*_*A*_ from expiratory flow rates *Φ*_*0*_ using gas density ρ and viscosity *µ*, and pleural pressure *P*_*pl*_.

#### From experimental MEFV with air to simulated MEFV with heliox

We assume that *P*_*pl*_ and *P*_*A*_ during MEFV maneuvers are dependent on respiratory muscle effort and independent from gas properties. Thus, similar alveolar pressure should be found during a maximal forced expiration with both air and heliox (27).

First, the pleural pressure *P*_*pl*_ is estimated from lung volume using the respiratory system pressure-volume relationship of Rahn (28) and of Agostoni and Hyatt (29) (A-box in **Figure 1**). These relationships have been determined in accordance with physiological measurements. Next, the alveolar pressure P_A_ is estimated by running the Stephano and Mauroy model on the MEFV data with air (S-box with *ρ*_*air*_ and *µ*_*air*_ in **Figure 1**).

Finally, to simulate expiratory flow with heliox, we used an iterative algorithm to determine which heliox flow rate was corresponding to the alveolar pressure *P*_*A*_ estimated with air (dotted-line box in **Figure 1**). At each iteration *i*, we run the Stephano and Mauroy model with the tested Heliox flow rate 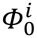 and the Heliox density and viscosity (S-box with *ρ*_*He*_ and *µ*_*He*_ in **Figure 1**) to compute the corresponding alveolar pressure 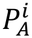. The heliox flow rate 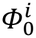 was adjusted to 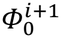 until 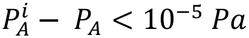 using the secant method (Correction-box, in **Figure 1**). The two initial flow values at the trachea needed for the secant method are the trivial no-flow value 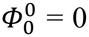 and the air flow rate measured experimentally that leads to the estimate of *P*_*A*_, i.e. 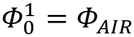.

### Statistical analysis

Normal distribution was evaluated using Shapiro-Wilk test and variance homogeneity was calculated using Levene test. FEV_1_ was compared between AIR and HeO_2_ using a paired Student t-test. Measured (AIR and HeO_2_) and simulated (SIMU, SIMU15, SIMU30) volume and flow rates were compared using a one-way repeated measure analysis of variance (ANOVA) and when a significant difference was found, multiple comparisons analysis was performed using the Tukey’s post-hoc test. Slope ratio was compared using two-way repeated measure ANOVA with two within-subjects variables (condition: AIR vs. HeO_2_ vs. SIMU vs. SIMU15 vs. SIMU30; volume: 20 to 80%FVC) and a Tukey post-hoc test was performed to determine where differences resided. Statistical analysis was performed using GraphPad Prism 10.5.0 (GraphPad Software, CA, USA). Data are presented as mean ± standard deviation (SD) unless stated otherwise and statistical significance was set at P < 0.05.

## RESULTS

All participants had spirometry data within the normal predictive values, and no participant showed values below the lower limit of normal (30) (**Table 1**). FEV_1_ (AIR: 3.22 ± 0.51 L vs. HeO_2_: 3.45 ± 0.49 L, P = 0.015), FEV_1_/FVC (AIR: 0.73 ± 0.08 vs. HeO_2_: 0.79 ± 0.07, P = 0.001) and FEF_25-75_ (AIR: 2.86 ± 0.96 L/s vs. HeO_2_: 3.92 ± 1.41 L/s, P < 0.001) were higher breathing HeO_2_ compared with air. Differences between measured and simulated spirometry variables are depicted in **Table 2**; measured and simulated MEFV curves are illustrated in **Figure 2**. The effect of heliox on expiratory flow was amplified when airway size was reduced, as indicated by the greater PEF, FEF_25_ and FEF_50_ in SIMU15 and SIMU30 compared with SIMU (**Table 2**). PEF_volume_ was 150 mL greater with HeO_2_ (**Table 2**). As the simulations were based on expiratory flow-volume data obtained with breathing air, the 150 mL difference in PEF_volume_ was retained when comparing HeO_2_ with SIMU, SIMU15, and SIMU30 (**Figure 2**). Expiratory flow-volume curves of each condition were aligned at PEF_volume_ with air, which reduced the difference in expiratory flow at a given lung volume between measured and simulated data (**Figure 3**). As indicated by the non-significant main effect of condition, the curvilinearity of MEFV curves was not different between conditions and between experimental and simulated data (**Figure 4**).

**Table 2.** Measured and simulated spirometry data.

| **Variable** | **AIR**<br>**(n = 14)** | **HeO<sub>2</sub>**<br>**(n = 14)** | **SIMU**<br>**(n = 14)** | **SIMU15**<br>**(n = 14)** | **SIMU30**<br>**(n = 14)** | **Condition**<br>**P value** |
|---|---|---|---|---|---|---|
| **FVC (L)** | 4.44 ± 0.71 | 4.38 ± 0.66 | 4.44 ± 0.71 | 4.44 ± 0.71 | 4.44 ± 0.71 | 0.307 |
| **PEF (L/s)** | 9.73 ± 1.82<sup>b,c,d,e</sup> | 13.48 ± 3.43 | 11.79 ± 3.93<sup>d,e</sup> | 12.19 ± 4.06<sup>e</sup> | 12.78 ± 4.12 | **<0.001** |
| **PEF<sub>volume</sub>(L)** | 0.61 ± 0.22 | 0.76 ± 0.26<sup>a,c,d,e</sup> | 0.61 ± 0.22 | 0.61 ± 0.22 | 0.61 ± 0.22 | **<0.001** |
| **FEF<sub>25</sub> (L/s)** | 7.75 ± 1.99<sup>b,c,d,e</sup> | 11.60 ± 3.35<sup>a,c,d,e</sup> | 8.88 ± 2.95<sup>d,e</sup> | 9.18 ± 3.05<sup>e</sup> | 9.66 ± 3.16 | **<0.001** |
| **FEF<sub>50</sub> (L/s)** | 3.72 ± 1.23<sup>b,c</sup> | 5.57 ± 1.89<sup>a,c,d,e</sup> | 3.48 ± 1.35<sup>d,e</sup> | 3.58 ± 1.42<sup>e</sup> | 3.73 ± 1.51 | **<0.001** |
| **FEF<sub>75</sub> (L/s)** | 0.97 ± 0.35 | 1.31 ± 0.70<sup>c,d,e</sup> | 0.92 ± 0.45 | 0.92 ± 0.46 | 0.93 ± 0.47 | **0.005** |

All values are expressed as means ± SD. FVC, forced vital capacity; FEF<sub>25</sub>, forced expiratory flow at 25% of FVC; FEF<sub>50</sub>, forced expiratory flow at 50% of FVC; FEF<sub>75</sub>, forced expiratory flow at 75% of FVC. <sup>a</sup>, P < 0.05 vs. AIR; <sup>b</sup>, P < 0.05 vs. HeO<sub>2</sub>; <sup>c</sup>, P < 0.05 vs. SIMU; <sup>d</sup>, P < 0.05 vs. SIMU15; <sup>e</sup>, P < 0.05 vs. SIMU30.

**Figure 2.**
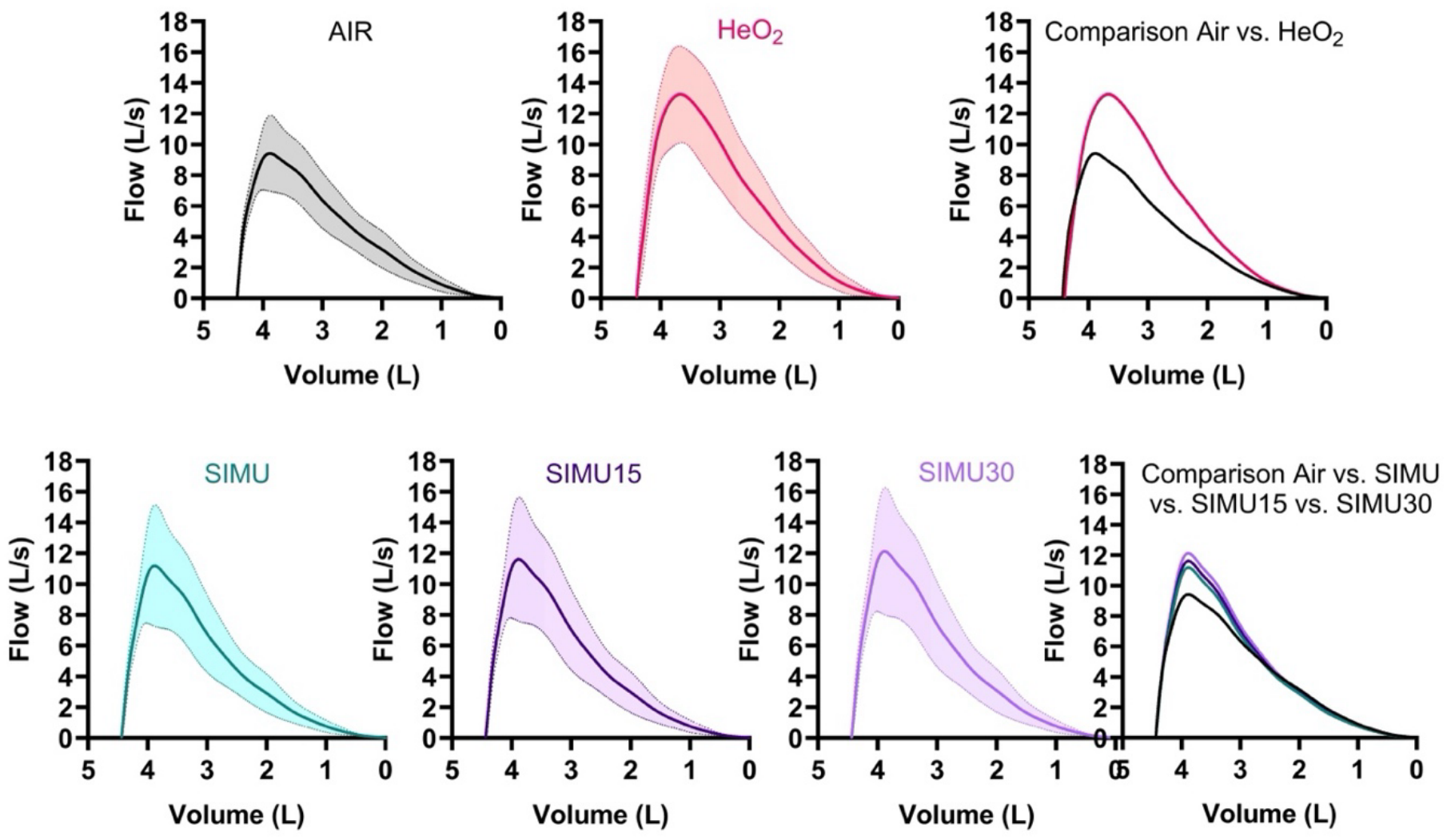
Maximal expiratory flow-volume curves measured with air (AIR) and heliox (HeO_2_), heliox simulated with no change in airway size (SIMU), simulated with a 15% (SIMU15) and a 30% (SIMU30) reduction in airway size. Data are presented as mean ± SD.

**Figure 3.**
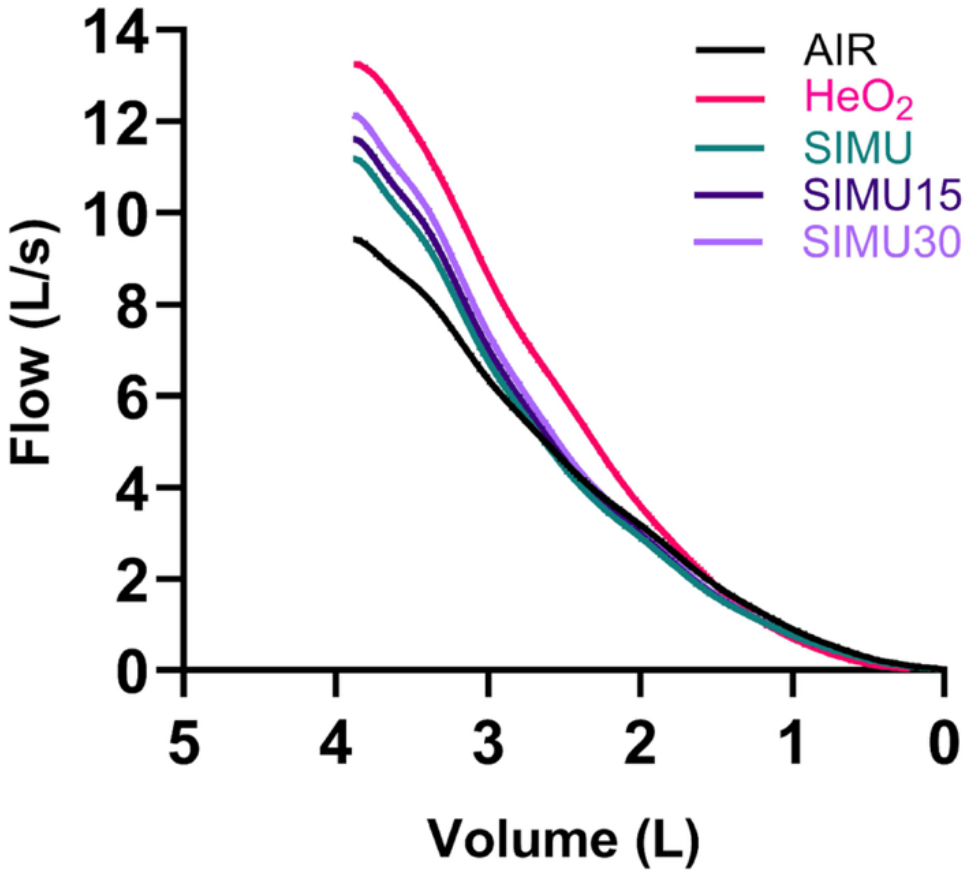
Maximal expiratory flow-volume curves aligned at peak expiratory flow volume (PEF_**volume**_**), measured with air (AIR) and heliox (HeO**_**2**_), and simulated with no change (SIMU), 15% (SIMU15) and 30% (SIMU30) reduction in airway size.

**Figure 4.**
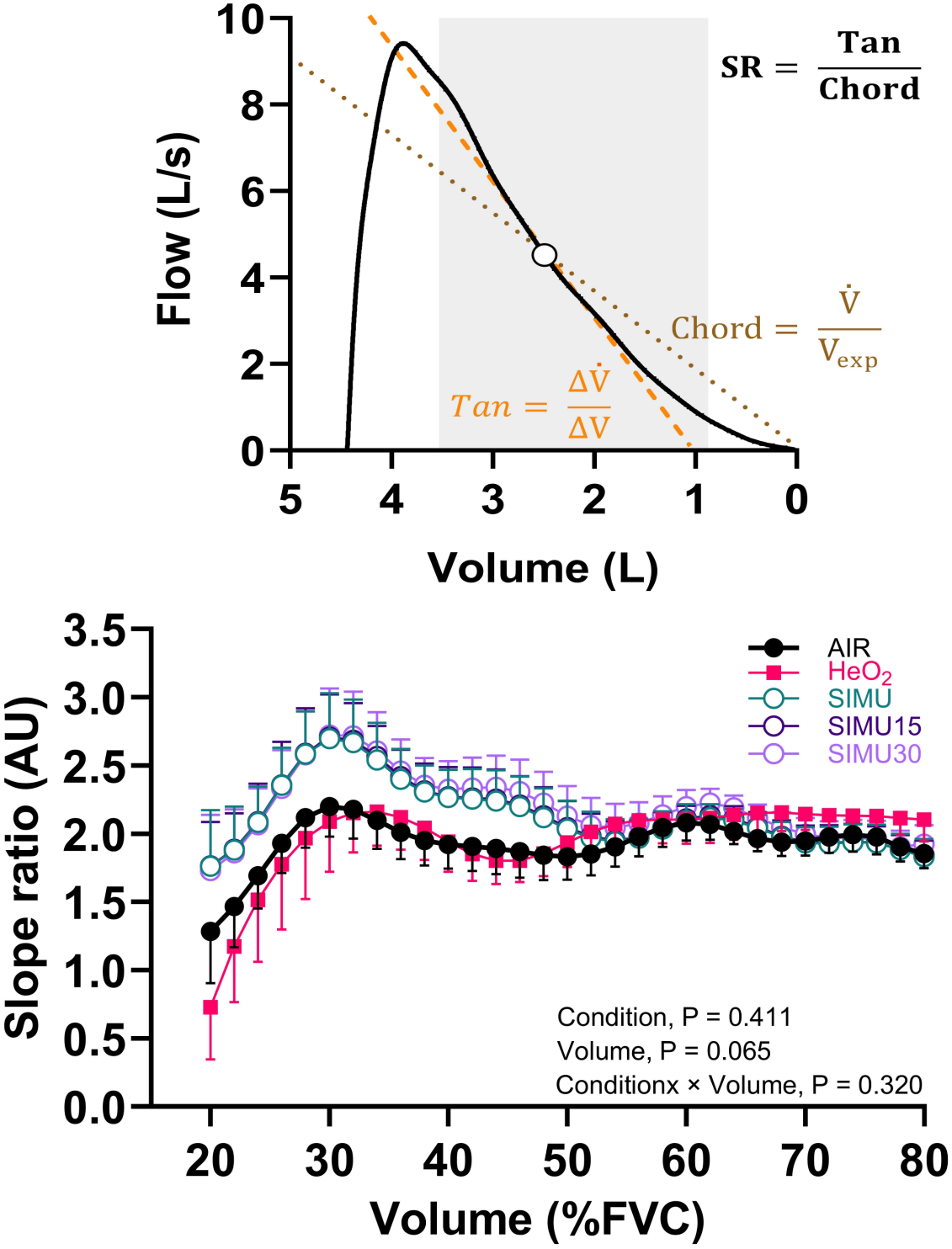
Slope ratio (SR) of maximal expiratory flow-volume (MEFV) curves. Representative illustration of the procedure used to determine the SR of a MEFV curve at one point of interest (open circle, upper panel). The grey area indicates the region of interest between 20% and 80% of FVC; the orange dashed line represents the tangent, and the brown dotted line represents the chord. SR values across the 30 points of interest (lower panel) for curves measured with air (AIR) and heliox (HeO_2_), and for simulations with no airway-size change (SIMU), a 15% reduction (SIMU15), and a 30% reduction (SIMU30). Data are presented as mean ± SEM.

At flow rates at the trachea of 5, 10 and 15 L/s, *R*_*e,n*_ is lower across the 17 generations of our pulmonary tree model when breathing heliox compared with air (**Figure 5**). When considering airway cross-sectional area from Lambert (17) and a flow of 1 L/s at the trachea, flow is laminar (*R*_*e,n*_< 2000) from the 5^th^ generation to the 16^th^ when breathing air and from the 2^nd^ generation to the 16^th^ when breathing HeO_2_. For a 15 L/s flow at the trachea, flow is laminar at the 12^th^ generation breathing air and from the 9^th^ generation breathing HeO_2_. On average, across the 17 generations studied, air flow is laminar across 3.1 ± 0.3 more generations compared with HeO_2_ flow.

**Figure 5.**
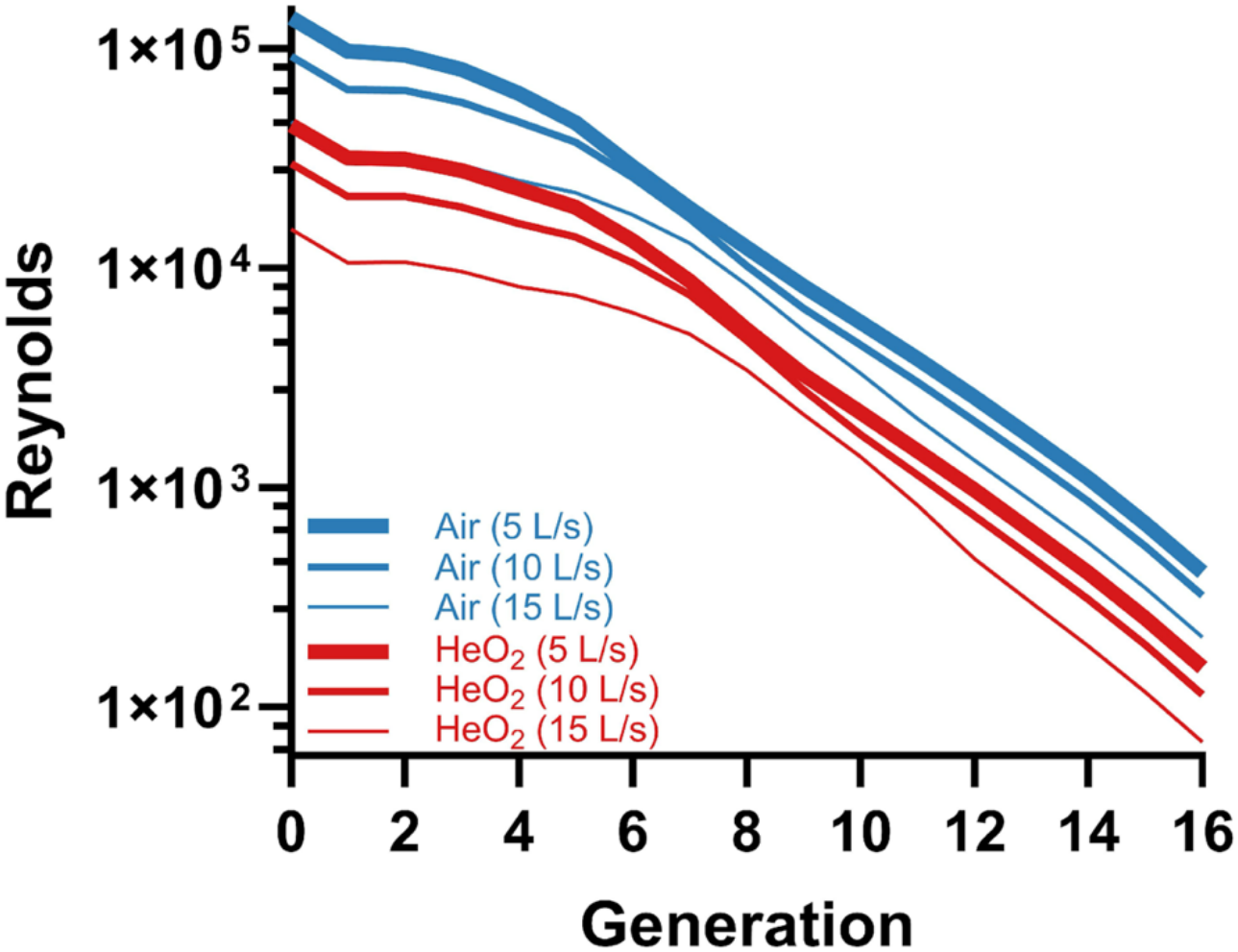
Reynolds number *R*_*e*,n_ versus airway generation index *n*, with air and HeO_2_ across lung generations for a flow rate at the trachea (generation 0) of 5, 10 and 15 L/s. Inertia effects become noticeable when *R*_*e*_ > 300 and they become dominant when *R*_*e*_ > 600. Heliox enhances expiratory flow gain as the Reynolds number increases. Turbulence might play a significant role when *R*_e_ > 2000.

The ratio of airway resistance with air (RES_AIR_) to airway resistance with heliox (RES_HeO2_) along each generation for a flow rate ranging from 1 to 15 L/s is depicted in **Figure 5**. For an expiratory flow rate of 1 L/s at the trachea, RES_HeO2_ becomes higher than RES_AIR_ from the 10^th^ generation. When flow rates increase, RES_HeO2_ becomes higher than RES_AIR_ deeper in the airways as it is the case for a flow rate of 14 L/s where 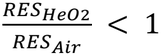 is found from the 16^th^ generation. In our model, when the regime of the flow shifts to Poiseuille (*R*_*e,n*_ < 600) and non-linear effects stop, the resistance ratio 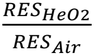 clips to the ratio of viscosity 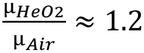. This occurred from the 10^th^ generation at 1 L/s, from the 14^th^ generation for 5 L/s and from the 16^th^ generation for 15 L/s (**Figure 6**).

**Figure 6.**
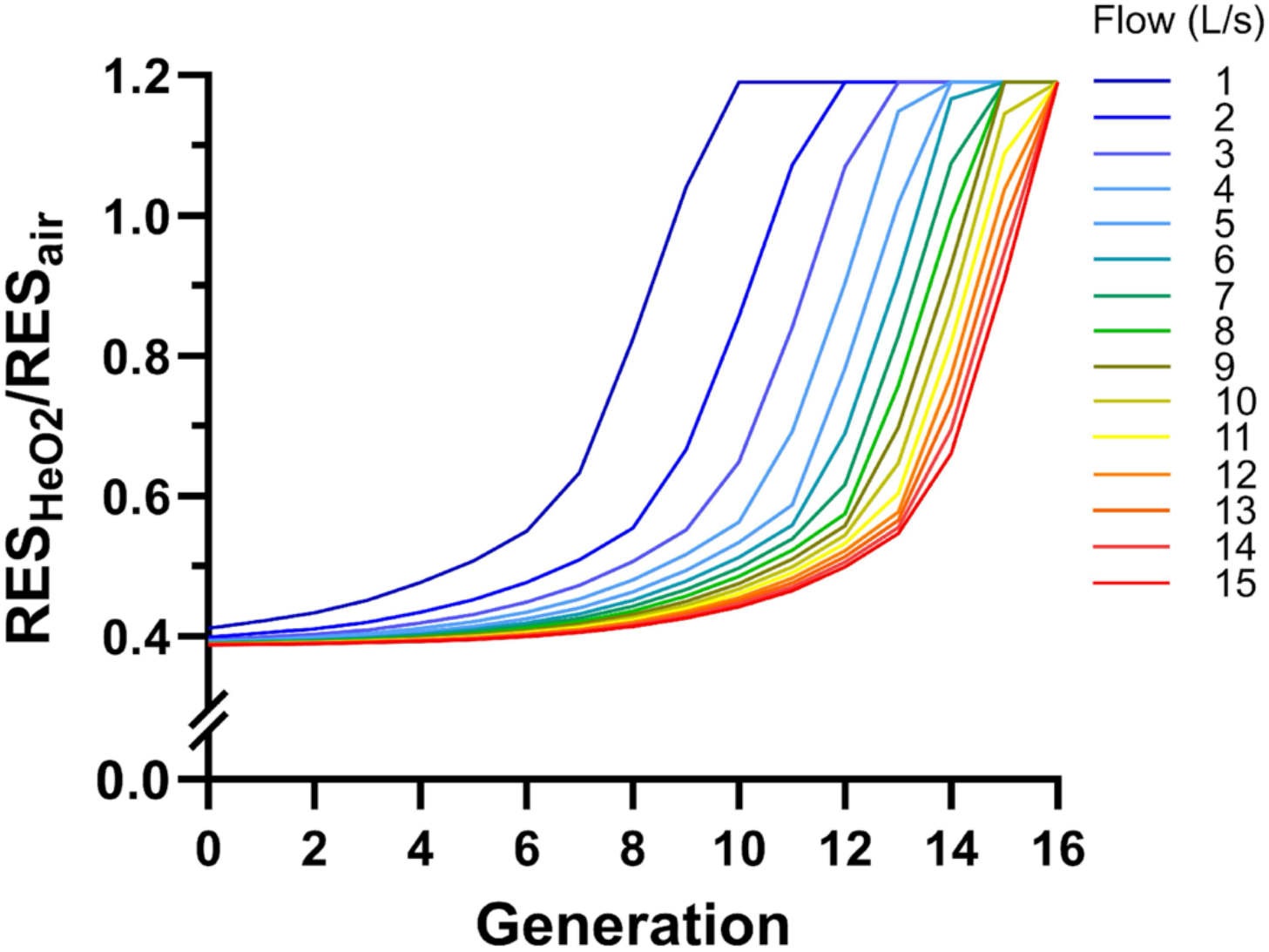
Ratio of air resistance by HeO_2_ resistance across lung generations for a flow from 1 to 15 L/s. A ratio 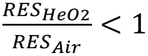 indicates that RES_HeO2_ is lower than RES_AIR_. When the ratio exceeds 1, resistance increases with heliox resulting in reduced benefits on expiratory flow. 1.19 corresponds to the viscosity ratio.

## DISCUSSION

The present study combines spirometry assessments and computational modeling to isolate the effects of gas density and viscosity on expiratory flow in master athletes, a model of successful aging. Breathing heliox increases expiratory flow at higher lung volumes without affecting flow at lower lung volumes or FVC (**Figure 2** and **Table 2**). These effects are primarily explained by regional differences in flow regime and airway resistance along the bronchial tree. In addition, our modeling approach indicates that age-related airway narrowing amplifies the beneficial effects of heliox on expiratory flow. This study thus provides a mechanistic framework to investigate how airway caliber influences expiratory flow responses to reduced gas density. By modifying airway dimensions independently to other mechanisms, the model helps isolate the contribution of airway geometry, which cannot be directly assessed experimentally. This approach helps interpret age-related changes in the expiratory flow response to heliox and improves our understanding of the mechanisms that limit expiratory flow in healthy aging while breathing air.

Pulmonary function data show that heliox breathing does not modify FVC, indicating that lowering gas density does not influence static lung volumes in master athletes which aligns with previous findings in younger adults (26). In contrast, heliox increased PEF, FEF_25_, and FEF_50_, while FEF_75_ remained unchanged. This pattern indicates that the effect of heliox is limited to the early phase of expiration, with no measurable benefit at low lung volumes. These findings indicate that reduced gas density enhances expiratory flow in regions where flow is highest, while having minimal impact in the distal lung where low Reynolds numbers are driven by viscosity.

Quantification of the MEFV curve shape using the slope ratio provides complementary insight into expiratory flow dynamics. Because expiratory flow between 20 and 80% of FVC is largely effort-independent (22), the absence of change in SR between air and HeO_2_ indicates that heliox does not alter airway emptying (21, 31). These findings are consistent with previous observations in younger adults and suggest that, in healthy aging, changes in gas physical properties do not substantially influence the overall distribution of airway resistance (21). Moreover, the similarity in SR between measured and simulated MEFV curves indicates that the model reproduced the overall curvilinearity of the experimental curves. Because airway narrowing was imposed uniformly across all airway generations in the simulations, the similarity between measured and simulated curves is consistent with a relatively uniform distribution of airway narrowing in our participants, rather than a markedly heterogeneous pattern or pronounced airway constriction (32).

The regional nature of the heliox effect reflects fundamental differences in airway resistance mechanisms along the bronchial tree. Airway resistance depends on both gas density and viscosity, but their relative contributions vary with flow conditions (15). In proximal airways, where flow velocities are high, inertial effects dominate and resistance is strongly influenced by gas density. In distal airways, where flow is low, resistance follows Poiseuille’s law and depends almost exclusively on viscosity. Accordingly, reducing gas density with heliox decreases resistance whereas its higher viscosity limits any benefit in distal airways (13–15). Our computational model provides mechanistic insight by isolating the effects of gas physical properties while maintaining constant lung geometry and driving pressure. Simulations show that the relative contribution of density- and viscosity-dependent resistance varies across airway generations and depends on flow magnitude. At higher tracheal flows, inertial effects extend deeper into the bronchial tree, whereas at lower flows, the transition to viscosity-dominated behavior occurs earlier. Importantly, because of its 2.6 times lower gas density at 37°C, heliox shifts this transition proximally. In contrast, in the most distal airways where there is a Poiseuille flow, resistance is determined exclusively by viscosity, and the slightly higher viscosity of heliox reduces expiratory flow, as also observed by others (26). Overall, these results indicate that the effect of heliox reflects a balance between reduced density-dependent resistance and increased viscous resistance, which varies across airway generations. By selectively reducing density-dependent resistance, we used heliox as a functional probe to investigate the distribution of airway resistance along the bronchial tree. When used in our model, it helps identify the airway generations in which reducing gas density most effectively decreases resistance and improves expiratory flow. This provides insight into how age-related changes in airway caliber and geometry may contribute to expiratory flow limitation breathing air, particularly during periods of elevated ventilation, such as during exercise. Analysis of Reynolds number provides further information. Regardless of gas composition, Reynolds number decreases along the bronchial tree as airway diameter and flow decline and is lower with heliox due to its lower density. This promotes a transition to laminar flow (i.e., Poiseuille flow) at a more proximal airway generation. While this shift increases the contribution of viscous resistance in more proximal regions, it simultaneously reduces inertial losses where flow velocities are highest, particularly at airway bifurcations. The reduction in density-dependent resistance outweighs the increase in viscous resistance under these conditions, resulting in a higher expiratory flow, induced by lower resistance of proximal airways. This explains why the effects of heliox are not uniform across the lung and is consistent with previous reports of reduced pressure losses during heliox breathing (15, 16, 26).

To investigate specifically the influence of aging-related structural changes, we simulated reductions in airway lumen area of 15% and 30% (4, 24). This approach allows isolating the influence of airway size as an independent variable, which is not feasible in vivo. Decreasing airway caliber increases overall resistance and amplifies the effect of heliox on expiratory flow for a given driving pressure. As shown by our model, for a given experimentally measured expiratory flow with air, narrower airways require a higher driving pressure to overcome their greater resistance. When the same driving pressure is then applied to the model using heliox properties, the lower gas density results in a greater predicted expiratory flow. These findings indicate that the benefits of heliox are more pronounced in smaller airways and should therefore increase expiratory flow more in older compared with younger adults for a given driving pressure. This highlights the greater limitation to generate expiratory flow in older adults, driven by increased airway resistance from age-related airway narrowing (5, 33). In contrast, separate simulations modifying airway compliance produce minimal changes in flow, suggesting that airway geometry, rather than elasticity, is the primary determinant of the observed differences.

The agreement between experimental and modeled MEFV curve shapes, as shown by the similar SR, suggests a consistent airway narrowing across airway generations. However, aging is a multifactorial process, and it should be noted that the mechanisms isolated in this study represent only a subset of the determinants of expiratory flow. Additionally, the use of a single, idealized lung geometry and simplified physics in the model may not capture all the biomechanical complexity or inter-individual variability, which could contribute to differences between simulated and measured data. These limitations should be considered in the context of the primary objective of the model, which was to investigate the relative influence of gas properties and airway geometry on expiratory flow rather than to provide an exact quantitative prediction of individual MEFV curves. Despite these limitations, the model offers a useful framework for testing mechanistic hypotheses and identifying how specific changes in airway structure may alter the distribution of flow resistance throughout the bronchial tree.

## CONCLUSION

Breathing a low-density gas mixture decreases expiratory flow resistance primarily in proximalairways by reducing density-dependent resistance under conditions where inertial effects dominate. This shifts the transition to laminar flow in more a proximal part of the bronchial tree and results in region-specific improvements in flow. These effects are amplified in the presence of smaller airway calibers, as observed with aging. The integration of physiological measurements with computational modeling provides a mechanistic framework for understanding how gas properties and airway structure interact to determine expiratory flow in the aging lung.

## ACKNOWLEDGEMENTS & GRANTS

We thank the participants for their time and effort dedicated to the participation in this study. Financial support was provided by Région Provence-Alpes-Côte d’Azur and the initiative of excellence (ANR-15-IDEX-01), joint, excellent, and dynamic initiative (JEDI) of the Université Côte d’Azur (UCA).

